# Influence of Nesting Material Composition on Tick Tube Use by *Peromyscus leucopus*

**DOI:** 10.1101/2022.01.07.475426

**Authors:** Jordan T. Mandli, Sydney E. Ring, Susan M. Paskewitz

## Abstract

Host-targeted acaricides are a valuable tool for the reduction of ticks and tick-borne disease. Tick tubes (also known as tick control tubes) are commercially available products containing permethrin-treated nesting materials. Through superficial acaricide application to *Peromyscus* mice, tick tubes reduce populations of the blacklegged tick, *Ixodes scapularis* Say. Results of prior field trials have varied, suggesting that mouse behavior as well as the scale of the intervention and the composition of the local host community are important determinants of efficacy. Here we evaluated behaviors related to nest material collection by *P. leucopus*. Two forms of nest materials used in commercial tick tube products (cotton batting and balls) were assessed through side-by-side comparisons over a four-week period. We quantified cotton uptake by monitoring weekly changes in material weight and used video surveillance to categorize and assess mouse behaviors. The odds of cotton batting being taken from tubes was 2.14 times greater than cotton balls but the process was less efficient; mice removed 0.35 g less cotton batting for each removal event and required 2.17 times longer to complete the removal. While cotton balls were readily carried in the jaws of mice, batting required separating smaller fragments from the mass before placement in the oral cavity. Video surveillance suggested that a small number of mice were super users and responsible for 22% of the 119 visits in which material was removed. Combined, material weight loss and video-captured removal events improve our understanding of host usage of nest materials but also raise questions about dissemination of the material in nests of the local mouse community.

## 1. INTRODUCTION

In North America, the white-footed-mouse, *Peromyscus leucopus* Rafinesque, is an important reservoir host that maintains tick-borne pathogens like *Borrelia burgdorferi* sensu stricto, the causative agent of Lyme disease, as well as *Anaplasma phagocytophilum*, and *Babesia microti* (Barbour 2017). Often found in great abundance, the *P. leucopus* is an excellent host for juvenile *Ixodes scapularis* Say ticks and is a highly competent reservoirs for these pathogens (Levine et al. 1985, Brunner et al. 2008, Barbour 2017). Methodologies for combating pathogens that persist within a sylvatic cycle between wildlife and tick vectors often focus on vector control (Wilson et al. 2015). These approaches seek to reduce host-seeking nymphal stages of *I. scapularis*, because this stage is responsible for a majority of the pathogen transmission to humans (Eisen et al. 2017). These nymphs come as the result of successful larval blood meals during the previous year. Strategies like yard treatments with insecticide sprays and manipulation of wild animal hosts are effective in reducing the abundance of *B. burgdorferi*-infected nymphs, although ecological variability, environmental concerns, and socioeconomic considerations have limited their usefulness on the community-level scale (Eisen and Dolan 2016, Eisen and Stafford 2020).

Host-targeted acaricides provide a directed approach that can be used on individual properties. These products are meant to reduce the risk of tick-borne disease (TBD) by two mechanisms: 1) reduction in tick abundance through direct mortality of juvenile ticks while on the host; and 2) the inhibition of pathogen transmission between infected and naïve tick cohorts by killing any infected ticks before they transmit pathogens to reservoir hosts. One such method that is commercially available is the “tick tube”, which consists of permethrin-treated cotton deployed in a cardboard applicator tube. Tubes are placed in preferred rodent habitat and mice gather cotton for use in the construction of their nests (Spielman 1988). Utilization of the permethrin-treated nesting material transfers a topical protective dose of acaricide to mice.

Initial studies evaluating the effectiveness of tick tubes on *I. scapularis* reported a substantial decline in both the number of infesting ticks on mice and the overall abundance of infected host-seeking *I. scapularis* in the year following deployment (Mather et al. 1988, Deblinger and Rimmer 1991). Recently, a 5-year small-scale study of homemade tick tubes revealed significant reductions in *B. burgdorferi*-infected questing nymph densities (66%) and questing nymph densities (54%) (Mandli et al. 2021). While these observations provide evidence for tick tube effectiveness, several trials in the northeastern United States did not result in similar reductions (Daniels et al. 1991, Stafford 1991, 1992, Ginsberg 1992, Jordan and Schulze 2019). Several of these studies documented a lack of treatment effectiveness following a period of low nesting material collection (Daniels et al. 1991, Stafford 1991, 1992, Ginsberg 1992). In some of these studies, periods of reduction in juvenile tick burdens on *P. leucopus* were observed, suggesting product efficacy, but this did not translate to reductions in questing nymphal densities in the following year. Differences in rodent collection and use of permethrin-treated material may be one of several factors (including scale of application and host community composition) responsible for these dissimilarities.

To examine mouse interactions with and use of nest material, and to determine whether cotton composition influences cotton retrieval, we examined the material usage, frequency of collection, and timing of mouse interactions. We compared two cotton nest material types used in commercially available tick tube products sold under the brands Damminix® (ECOHEALTH, INC., Brookline, MA) and Thermacell® (Thermacell Repellents, Inc., Bedford, MA). These products share a similar design and permethrin formulation but differ in the cotton substrate. Given the similarity between products, we hypothesized that there would be no difference in material acceptance and removal.

## 2. MATERIALS AND METHODS

### 2.1 Study location

The study was conducted at the University of Wisconsin Arboretum located in Madison, WI (43.047764, -89.422732). Experiments were implemented in a maple-dominated forest with coarse woody debris and limited underbrush. Previous trapping efforts indicated a small mammal population consisting of white-footed mice, eastern chipmunk (*Tamias striatus* Linnaeus), eastern grey squirrel (*Sciurus carolinensis* Gmelin), southern flying squirrel (*Glaucomys volans* Linnaeus), and northern short-tailed shrew (*Blarina brevicauda* Say).

### 2.2 Experimental design

Thirty pairs of commercial tick tubes containing untreated material, provided by Thermacell Inc. (Bedford, MA), were staked down 10 meters apart along a single 290 meter transect. Each pair comprised a tube containing cotton batting and another with cotton balls (Figure 1). Tick tubes were swapped with a second set of paired tubes at weekly intervals over four weeks (7/12/18 – 8/9/18) to facilitate tube measurements. The first set (set A) was deployed on odd weeks (week 1 and 3) and the second set (set B) was deployed on even weeks (week 2 and 4). All tubes were weighed at the beginning of the trial on an analytical scale (Mettler-Toledo LLC, Columbus, OH) and cotton was re-distributed to equilibrate weights. At the end of each succeeding week, the entire tick tube including the contents was weighed (whole-tube). Debris, slugs, and dirt were manually removed from tubes to minimize confounding. Because we saw unexpected increases in tube weight in weeks 1 and 2, we separated the cotton from the tube during weeks 3 and 4 and weighed it (cotton-only), before reloading the tubes for return to the field. Tick tubes were reused between weeks (odd or even) except in the event that > 80% of cotton was removed or the tube was damaged. In such instances, the tick tube was replaced with a new one. Tube disturbance and abandoned cotton around the immediate area were recorded and photographs taken for a visual record. Cotton that was observed outside of tubes was left undisturbed.

**Figure 1:**
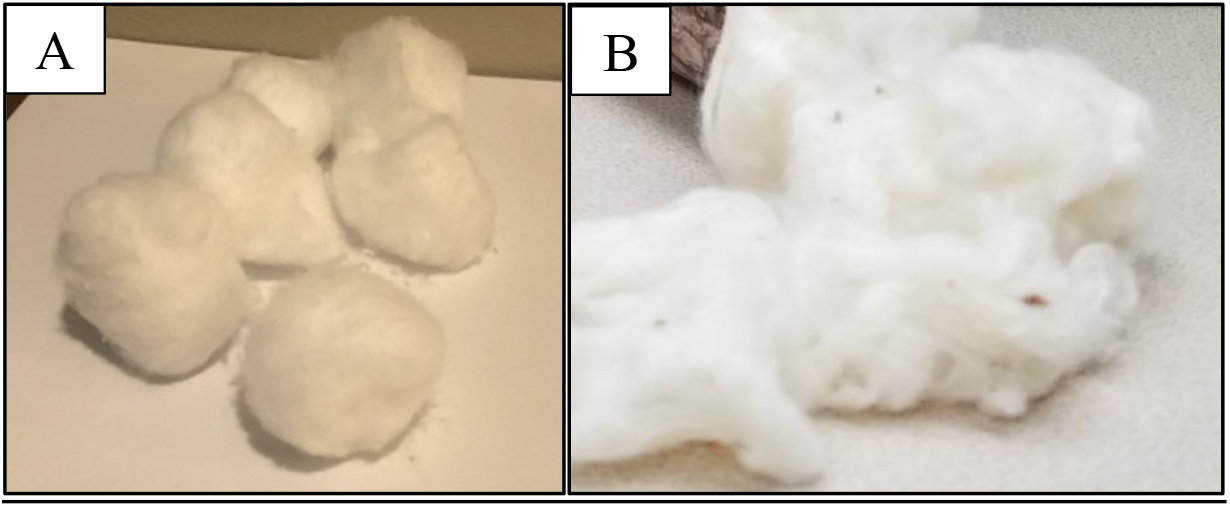
Depiction of material substrates. A) Cotton balls and B) Cotton batting.

### 2.3 Camera trap video collection

Four camera trap systems were deployed on 8/22/18 at 12:00 PM and placed along the transect in locations where tubes showed the greatest decrease in weight during paired trap weight comparisons. A fifth trap was placed shortly thereafter on 8/24/18 at 12:00 PM. Camera traps were spaced 10 to 50 meters apart and remained in the field for 6 to 8 consecutive days until recovery on 8/30/18 at 12:00 pm. Camera trap systems consisted of a Bushnell TM Essential E3 Trail camera (Bushnell Outdoor Products, Overland Park, KS) mounted to a PVC structure allowing for top-down visualization (Figure 2). Each camera trap offered an approximate one square-foot field of view. Tick tube pairs were centered under the camera trap apparatus. Video was downloaded every 24 hours at noon. Tubes were replaced every 48 hours and were returned to the lab for drying and weight measurement. Cameras began recording two seconds after motion was detected and were set to capture 15 seconds of footage (clip). Video was analyzed to reflect a single visit made by a mouse, and occasionally spanned multiple contiguous clips. Clips that featured two mice were evaluated as two separate visits, each referring to the specific activity of the individual mouse.

**Figure 2:**
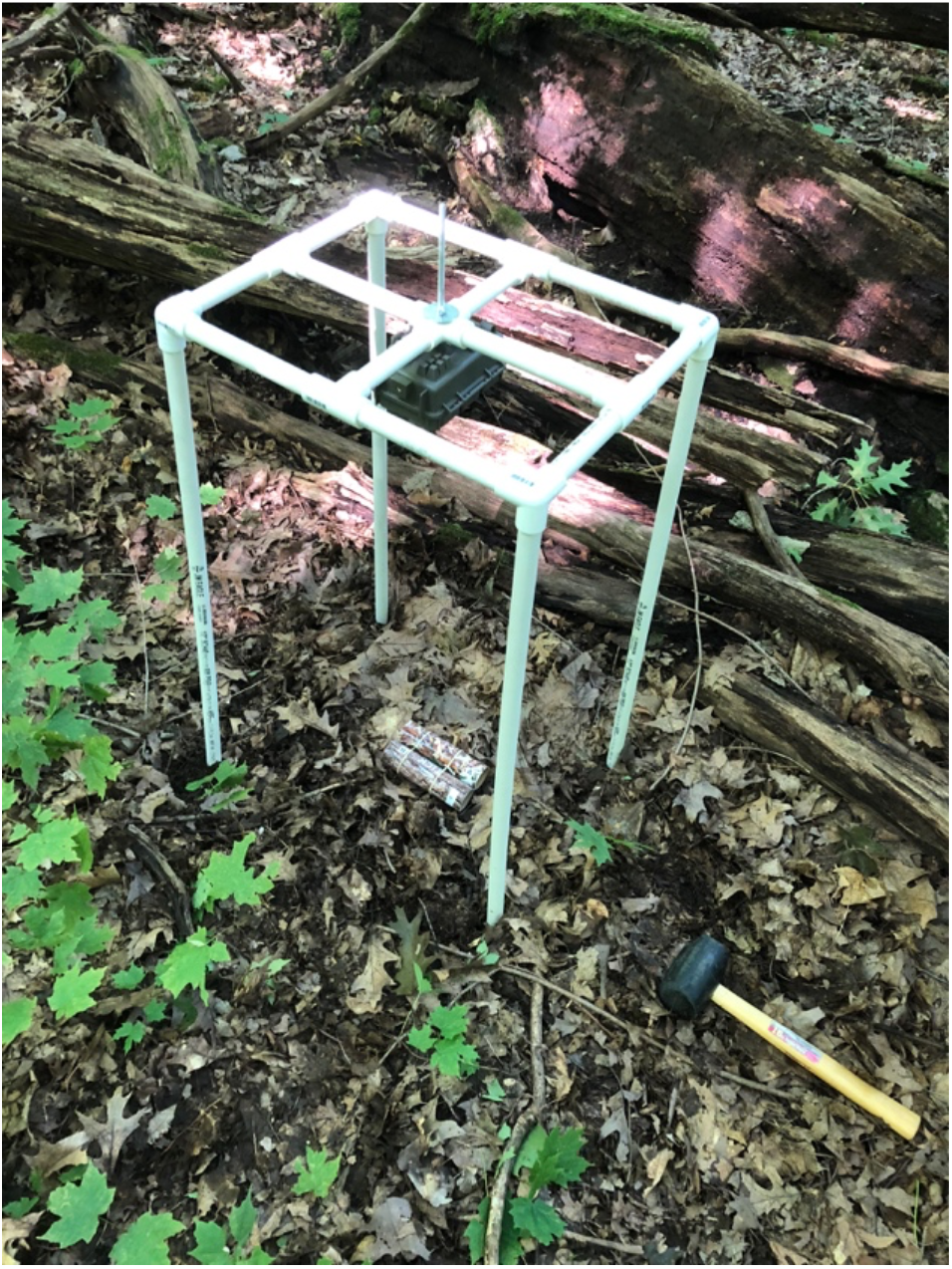
Mounted camera trap setup with tick tubes in the field

A coding key was created to quantify patterns of mouse activity. For analysis, outcomes were defined as follows: (i) ***visitation***, summarized as the number of visits to an individual tick tube per hour, regardless of collection outcome; (ii) ***removal outcome***, assessed whether nesting material was removed from the field of view during a single mouse visit and in the event that material was taken, which material. Removal outcome was scored as either unknown (UNK), no material taken (NONE), cotton balls taken (CBL), cotton batting taken (CBT), or both nesting materials taken (CBL+CBT); (iii) ***time in frame***, time (sec) in which a single mouse remains in camera field of view; (iv) ***time period***, hour of mouse visit was grouped into four three-hour time periods identified as sunset (SS = 19:00-22:00), late evening (LE = 22:00-1:00), early morning (EM = 1:00-4:00), and sunrise (SR = 4:00-7:00).

### 2.4 Statistical analysis

To evaluate the potential of confounding due to water and debris accumulation on whole-tube weights, a Pearson correlation test was used to examine the relationship between cotton-only weight changes and corresponding whole-tube weight changes during weeks 3 and 4. Follow-up robust linear regression was used to estimate the discrepancy between whole-tube and cotton-only weight changes. Welch’s t-tests were used to detect differences in whole-tube and cotton-only weight changes for each individual week when appropriate and for the cumulative two or four-week period (cotton-only = Weeks 3 and 4 only).

To assess collection behaviors captured by video, differences in the probability of observed removal outcomes for each nest material type were analyzed by chi-square analysis with a Yates’ continuity correction. Correlation between material weight changes and removal outcomes during camera trapping were assessed using a Pearson correlation test. Efficiency of removal by material composition was subsequently estimated by robust linear regression.

Analysis of visitation and time in frame was conducted using general linear mixed effect models (GLMMs) fitted by maximum likelihood with Laplace Approximation and log-link function (negative-binomial distribution). Time period was included as the sole visitation explanatory variable. In a separate model, time period and removal outcome were included as time in frame explanatory variables. We used camera trap nested in date as a random effect in all models to account for pseudo-replication.

Model selection was carried out based on corrected Akaike Information Criterion for small sample sizes (AICc) (Burnham and Anderson 2002). The simplest biologically compelling model within 2 AICc of the model with the lowest AICc value was selected as the best model. Analyses were carried out in R, version 1.4.1717 (R Core Team 2021). Robust linear regression was competed using the lmrob functions of the robustbase package v.93-9. GLMMs were developed in glmer.nb functions of the lme4 package (Bates et al. 2015).

## 3. RESULTS

### 3.1 Head to head comparison

All tick tubes were successfully retrieved from the field at the end of each week and weighed back in the lab. Twenty-one tubes (18%) were replaced with a new one prior to redeployment due to low cotton reserves (< 80% of pre-deployment weight), substantial damage caused by animal activity, or tube structural instability (usually because the tube was too wet). In some cases, physical deformation inhibited access to nesting materials. At the end of the week tick tubes were occasionally sodden, caked in dirt, or covered in slugs or slug excretions. Despite our best efforts to clean and dry tubes before measurement, 54% (n = 240) whole tube weight changes increased during the study. During the particularly wet conditions of week one, 75% (n = 60) of whole tubes were heavier at the end of the week. Interestingly, robust linear regression showed a strong relationship between all cotton-only weight change (g) and whole-tube weight change (g) in weeks 3 and 4 [r(118) = 0.99, p-value < .0001]. The slope coefficient signified that whole-tube weights captured 95% of each gram of cotton-only weight loss thereby suggesting minimal discrepancy between measurements.

A significant difference in whole tube weight changes between materials was detected in Differences in whole-tube changes in weight between materials were significant in week one (t = 1.39, df = 29, p-value = 0.024), when cotton batting tubes had greater reductions, and in week 3 (t = 2.20, df = 29, p-value = 0.036) when cotton ball tubes had greater declines. Significantly greater weight changes were observed for cotton balls compared to cotton batting in week 3 (t = 2.47, df= 29, p-value = 0.020) and for the combined weeks 3 and 4 period (t = 2.31, df= 59, p-value = 0.024) (Figure 3). The largest cotton-only changes in weight were observed in week 4 with reductions of 33% (balls) and 27% (batting) (Figure 3).

**Figure 3:**
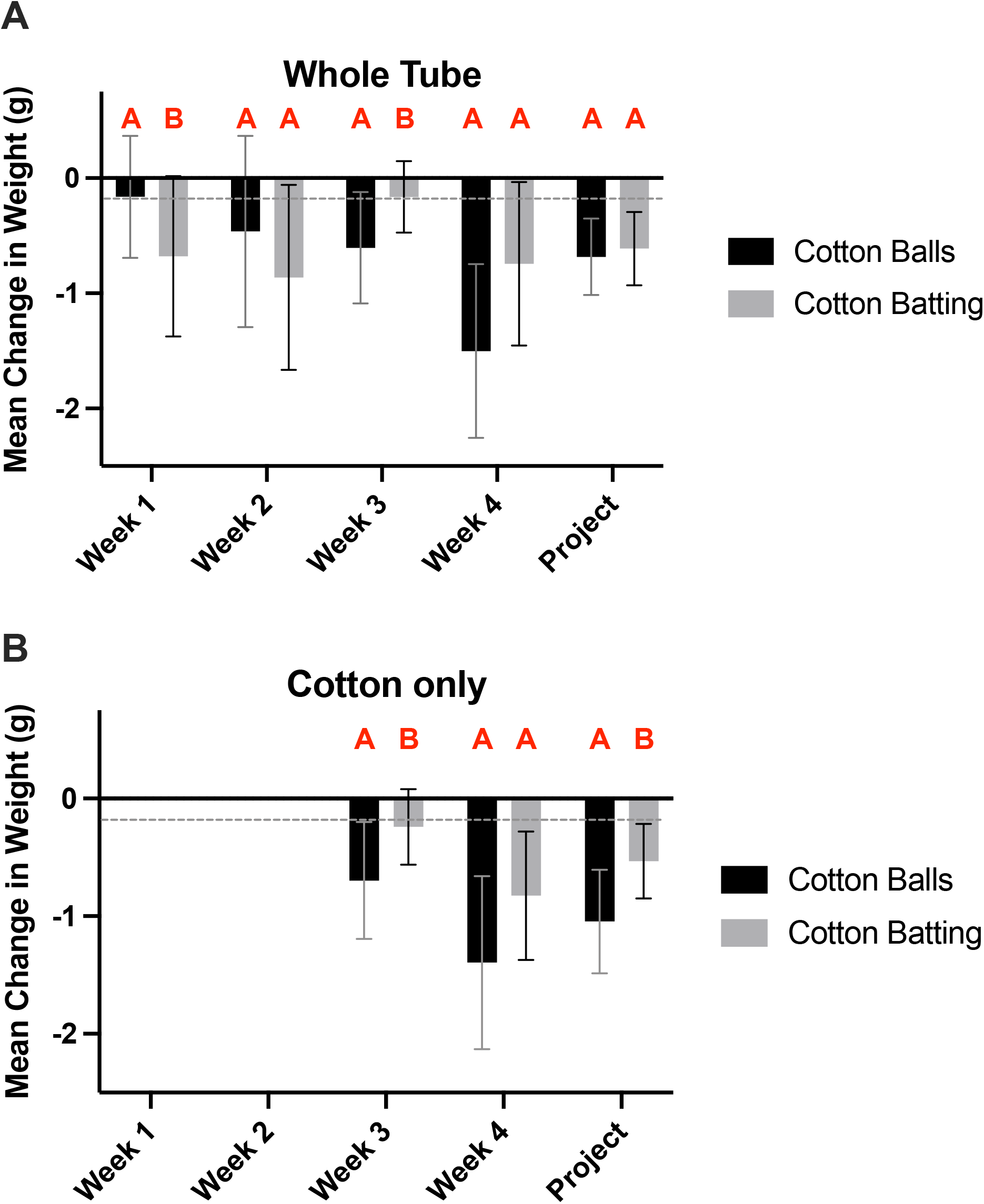
Mean weight change for whole-tube (A) and cotton-only (B) by material composition. Negative values indicate a loss in weight. Values are subdivided by week and a cumulative four-week period (Project). Paired bars with the same letter are not significantly different by paired t-test.

### 3.2 Camera trap evidence of animal activity

Cameras traps were functional during much of the observation period. A single camera failed to record footage between 8/24/18 to 8/26/18. Reduced clip times occasionally occurred, but simply increased the number of clips associated with a single visit. Collectively, 187 *P. leucopus* visits were captured during 864 total hours of observation (0.22 trips/hour). Footage of *S. carolinensis* was obtained on twenty occasions and a *Didelphis virginiana* (opossum) was observed once, but nesting materials were solely gathered by *P. leucopus*. The number of white-footed mouse visits varied depending on the camera trap (range: 0.14 - 0.29 trips/hour) and overnight time period (range: 0.08 – 0.39 trips/hour). Visitation occurred only after sundown between 20:00 to 5:00 with highest activity around 23:00 (Figure 4). Two spikes in visitation were detected at 22:00 and 4:00 but were largely attributed to specific camera traps on particular nights. Close examination of mouse features, size, and the close timing of visits suggested that individual mice were repeatedly returning to the tubes during spikes in activity. Of the 119 removal outcomes across the study, we suspect at least 22% (n=26) were the result of just three mice. Frequency of visits were explained by time period, with visitation negatively associated with the sunset (SS) and sunrise (SR) (Table 1).

**Table 1:**
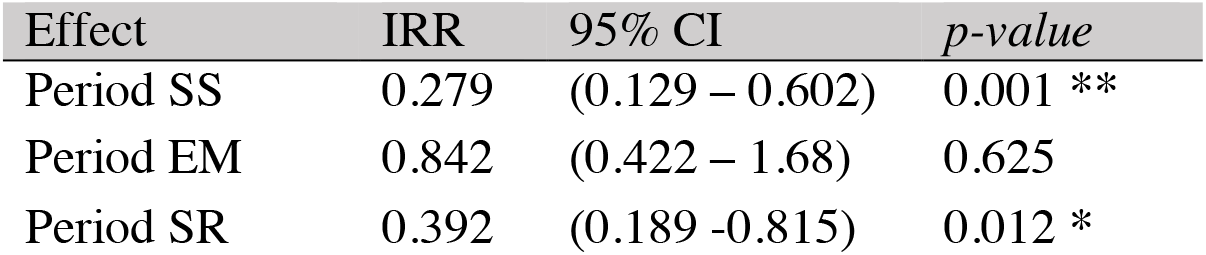
The instances of interaction between *P. leucopus* and tick tubes in relation to explanatory variable selected for the best model. Time period = four levels: four levels: sunset (SS), early morning (EM), sunrise (SR), and late evening (LE) as reference. Reported as incident rate ratios (IRR) with 95% confidence interval and p-value.

| Effect | IRR | 95% CI | p-value |
| --- | --- | --- | --- |
| Period SS | 0.279 | (0.129 – 0.602) | 0.001 ** |
| Period EM | 0.842 | (0.422 – 1.68) | 0.625 |
| Period SR | 0.392 | (0.189 -0.815) | 0.012 * |

**Figure 4:**
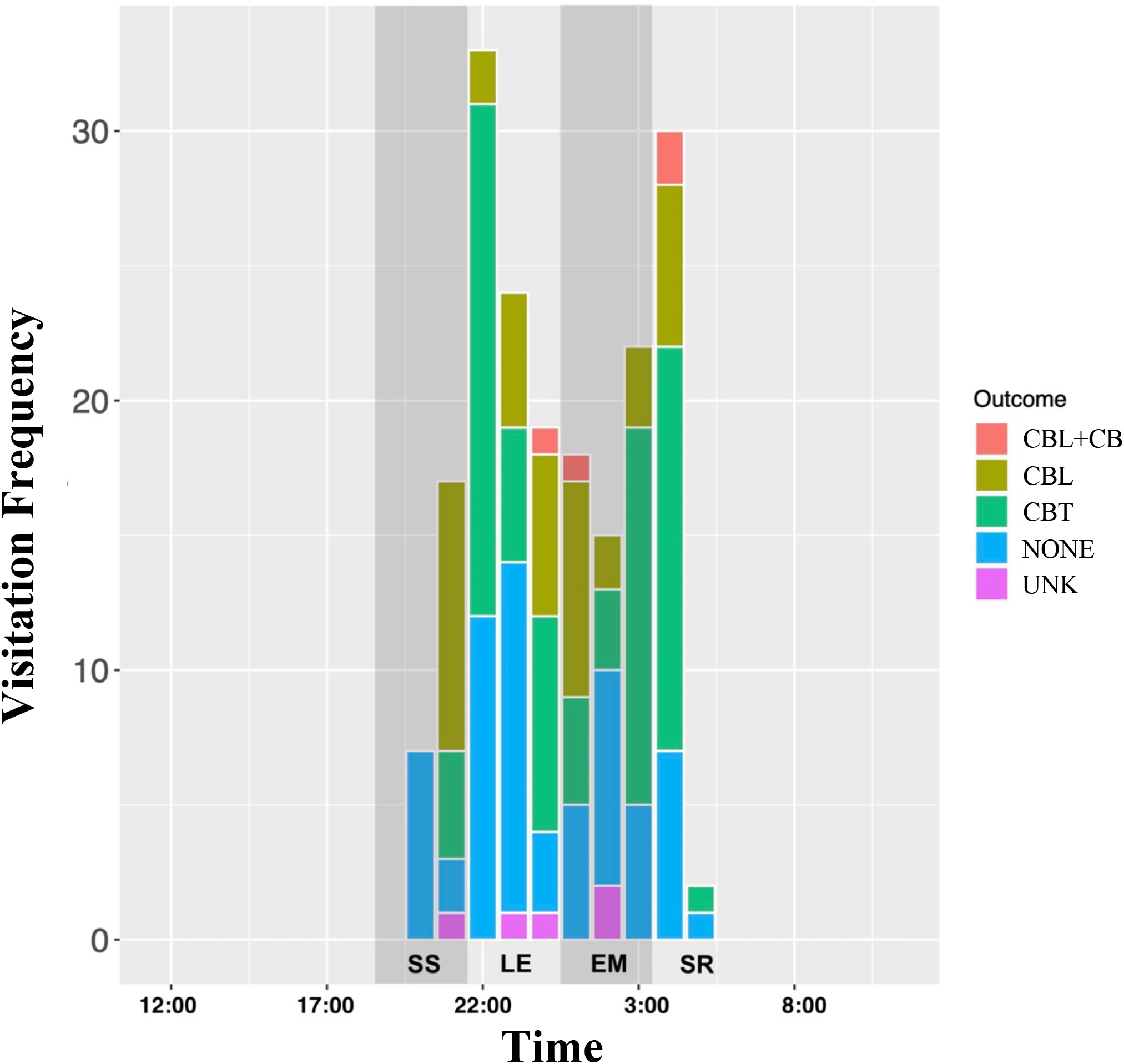
Visitation frequency by hour grouped by time periods sunset (SS), late evening (LE), early morning (EM), and sunrise (SR). Columns are subdivided by removal outcome; unknown outcome (UNK), no material taken (NONE), cotton batting taken (CBT), cotton balls taken (CBL), or both nesting materials taken (CBL+CBT).

### 3.3 Removal outcome

Of the 187 visits, we observed 63 instances of no material taken, 42 instances cotton balls taken, 73 instances of cotton batting taken, 4 instances of both cotton batting and balls taken, and 5 instances of unknown outcomes (Figure 4). Visits were predominately made by single mice but footage of two mice visiting a tube simultaneously was captured (see video clip 1). Mice carried cotton in their mouths for transport and were more likely to leave with cotton batting compared to cotton balls (OR = 2.14, *χ*^2^ = 15.1, df 1, p-value < 0.001). However, tube (t = 3.6, df = 18, p-value = 0.002) and cotton weight change (t= 3.7, df = 18, p-value = 0.001) measurements collected in parallel with video indicated less batting material was removed compared to balls. Simple linear regression showed a significant relationship between removal outcomes and weight changes for balls [r(17) = 0.60, p-value 0.001] and batting [r(17) = 0.83, p-value < 0.001]. Models predicted a reduction of 0.63 g per ball removal event and 0.28 g per batting removal event. A difference of 0.35 g per event. Delays in our camera trap system constrained our ability to detect rapid removal events (< 2 second). The extent of the deficiency remains unknown, but likely underestimates the speed and occurrence of cotton ball removal (see Clip 2).

### 3.4 Time in frame

As a proxy for time to collect material, we estimated the time each mouse remained in frame. Mice were observed for 1 to 96 seconds with an overall median time of 8.0 seconds (95% CI: 5.0 – 10). CBL+CBT outcomes demonstrated the longest times in frame (median = 60 sec, 95% CI: 30-80 sec, n = 4) followed by CBT (median = 14 sec, 95% CI: 12-16 sec, n = 73), CBL (median = 6 sec, 95% CI: 4.0-9.5 sec, n = 42), NONE (median = 4 sec, 95% CI: 3.0-5.0 sec, n = 63), and UNK (median = 1 sec, 95% CI = 1-11 sec, n = 5) (Figure 5). Time in frame was associated with time period and removal outcome, with longer visits occurring during sunset (SS) followed by sunrise (SR) and early morning (EM) compared to late evening (LE). Dual removal events (CBl + CBT) and CBT were associated with increased time in frame compared to NONE (Table 2). When comparing the time to remove individual treatments, CBT mice spent 2.17 times longer in frame compared to CBL.

**Table 2:** The time in frame of *P. leucopus* in relation to explanatory variables selected for the best models. Time period = four levels: sunset (SS), early morning (EM), sunrise (SR), and late evening (LE) as reference. Removal outcome = five levels: CBL+CBT, CBL, CBT, UNK, and NONE as reference. Reported as incident rate ratios (IRR) with 95% confidence interval and p-value.

| Effect: | IRR | 95% CI | p-value |
| --- | --- | --- | --- |
| Period SS | 1.91 | (1.19 – 3.10) | 0.008 ** |
| Period EM | 1.58 | (1.05 – 2.37) | 0.028 * |
| Period SR | 1.66 | (1.10 – 2.52) | 0.017 * |
| Material CBL+CBT | 6.53 | (2.86 – 17.4) | < 0.001 *** |
| Material CBL | 1.31 | (0.87 – 1.99) | 0.198 |
| Material CBT | 2.85 | (2.00 – 4.06) | < 0.001 *** |
| Material UNK | 0.452 | (0.18 – 1.21) | 0.098 |

**Figure 5:**
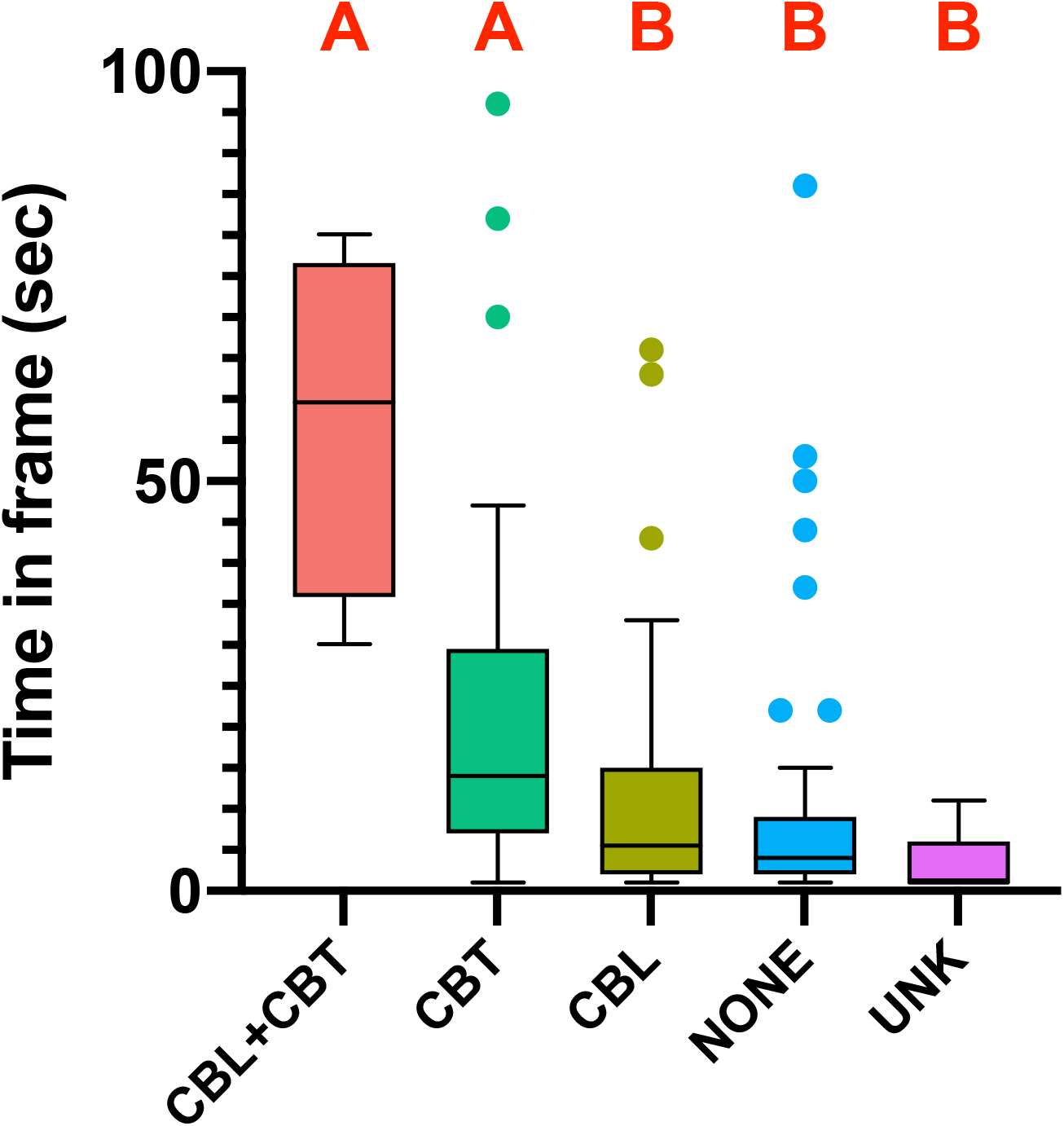
Median time in frame for *P. leucopus* by removal outcome. These include unknown outcome (UNK), no material taken (NONE), cotton balls taken (CBL), cotton batting taken (CBT), or both nesting materials taken (CBL+CBT). Boxplots with the same red letter (at top of graph) are not significantly different by Mann-Whitney Test.

### 3.5 Video observations

“Cotton dumping” occurred when nesting material was removed from the tube but was left within ∼3 meters of the tube (Figure 6). Almost all dumped material was eventually retrieved, presumably by mice. While cotton balls and cotton batting were dumped with nearly equal frequency, cotton batting was typically observed in much larger quantities. This was unsurprising given that mice were required to separate individual tufts of cotton batting from the larger wad to pack into their cheeks before carrying it off (see Clip 4). As such, there were multiple occurrences in which a majority of the cotton batting contents were carried off in a single visit (see Clip 3). Alternatively, cotton balls were gathered whole and held in the jaws of a mouse (see Clip 5). Mice sometimes were observed removing multiple balls or both materials in a single visit (see Clips 6 and 7).

**Figure 6:**
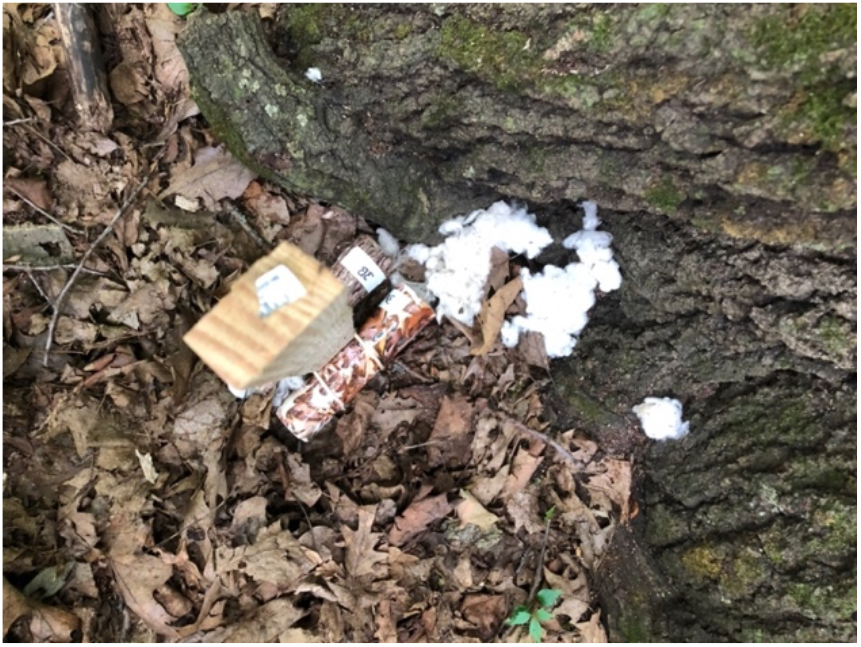
Example of cotton dumping event in which nesting material is deposited in tube surroundings.

## 4. DISCUSSION

As expected, white-footed mice were the sole consumers of cotton nest materials substrates within our study site. In the absence of an alternative cotton user, tick tube effectiveness depends on the relative importance of *P. leucopus* as a tick host and pathogen reservoir as well as rodent use of the product. This study demonstrates that cotton composition influences the removal of nesting material from tick tubes by *P. leucopus*. Cotton batting was removed much more frequently than cotton balls (e.g. 73 vs 42 removal events captured on video) but there were greater declines in weight for cotton balls compared to cotton batting both in the weeklong side-by-side comparisons and in the single night video capture experiments. Combining video surveillance of tick tube visits with nest material weight changes the following day, we predicted that approximately 0.63 g was removed per ball removal event and 0.28 g per batting removal event. Batting removal was also associated with prolonged time-in-frame and was more likely to occur during time periods just after sunset and before sunrise. Conversely, tick tube visitation increased during the darker hours of late evening and early morning but these visits predominately consisted of quick removal events (i.e. CBL and NONE) perhaps signifying greater perceived risk at these times.

Patterns of collection by *P. leucopus* varied depending on cotton composition due to the mechanisms that mice use to retrieve cotton. Mice separated individual tufts of cotton batting from the larger wad and packed these into their cheeks. This process was both time consuming and limited by mouse oral capacity resulting in longer time-in-frame and smaller reductions in batting weight. By contrast, cotton balls were gathered whole and held in the jaws of a mouse, which facilitated faster removal and transport. Efficiency of material collection may play an important role in cotton dissemination to nests. Modeling efficiencies under different population densities and mouse activity patterns (seasons) may reveal if one cotton material is better suited to exposing mice to permethrin.

In this study, single mice repeatedly collected large portions of nesting material. In particular, three mice were implicated in 22% of the total 119 removal events. This observation raises the question of how the behavior of individual mice might affect the efficacy of product dissemination among nests in an area. These high intensity periods also suggest that measuring decreases in weight or volume of material from tick tubes could be misleading as indicators of the likelihood of acaricide application to the local mouse community, if, for example, all or most of the cotton was disseminated to a single nest. Moreover, cotton dumping also introduces an additional disconnect between material removal from tick tubes and arrival to nests. Therefore, tick tube weight changes and percentage of cotton removal cannot be used as a proxy for treatment of the local *P. leucopus* population. Further research allowing for differentiation of mice (e.g., PIT tags (Godsall et al. 2014)) may better elucidate the portion of the population using tick tubes.

Research on the utilization of permethrin-treated nesting products by mice is lacking, thus critical factors influencing acaricide application to a population of mice are largely unknown. Some evidence exists supporting the possibility that mice that are not collecting cotton are still being treated in the nests (Ginsberg 1992). White-footed mice inhabit multiple nests over short periods of time and even share nests simultaneously with other (adult) mice (Wolff and Hurlbutt 1982, Madison et al. 1984, Larson et al. 2019). Therefore, incorporation of permethrin-treated cotton into a single nest by one mouse could treat many additional mice. Another important factor is the longevity of permethrin within the nest material. Ginsberg (1992) demonstrated stability of permethrin in soils surrounding treated cotton for up to 25 days. This provides an indirect measure of the minimum lifespan of treated cotton in the environment. Further study is needed to confirm these observations and to determine 1) how cotton is incorporated in the nest, 2) how many mice in a population are treated and over what timespan, 3) how much, where, and how often does permethrin need to be applied on mice to be protective, and 4) how mouse density influences dispersal of the cotton into nests. Recent studies indicating the presence of larval and nymphal blacklegged ticks in nests are also intriguing and could be especially relevant for tick tube use if some immature ticks are successfully overwintering in untreated nests and able to attach to hosts earlier in the season than those relying on questing in environments outside the nest (Larson et al. 2019). Radio telemetry and nesting boxes may offer insight into these questions and provide valuable information towards the refinement of tick tube deployment in varying locales (Madison et al. 1984, Larson et al. 2019).

Animal foraging behaviors are a balance of perceived risk versus reward in which prey demonstrate ani-predator behaviors under different risk stimuli (Lima and Bednekoff 1999). Thus, white-footed mice are likely to change their interactions with tick tubes depending on abiotic and biotic cues. During the hours closest to sunrise and sunset, tick tube visitation was low but time in frame was high. Later at night this pattern switches, with increased frequency but shorter visits observed. One explanation for the change in behavior could relate to perceived threats. Predator cues have been shown to influence foraging behaviors and are expected to decrease mouse interaction with tick tubes (Brown and Kotler 2004). Shorter visits in late evening and early morning hours could reflect a response to greater predator threats during these periods in comparison with the hours around sunset and sunrise. Given the prolonged collection time and the need for multiple visits associated with collecting amounts of batting equivalent to a single cotton ball, perceived predator threats may be more likely to affect mouse use of batting in comparison with balls. Further examination of mouse and tick tube interactions under varying predator signals may offer additional insight into the sources of variation in cotton nesting material collection that have been observed (Stafford 1991, 1992).

Ambient temperature has been shown to affect rodent activity within the temperate climate zone (Orrock and Danielson 2009). As the difference between body and ambient temperature grow, rodent metabolic rate increases to compensate for heat loss (Vickery and Bider 1981). Reduced temperature acts as a cue for *P. leucopus* to increase nesting behavior especially in populations from southern latitudes (Heath and Lynch 1983, Stafford 1992). Therefore, tick tube use is likely subject to variability due to seasonality in different geographical regions. Further study of seasonality would be valuable towards assessing patterns of collection in early spring prior to questing activity or late fall before mouse torpor in order to optimize dispersal of nesting materials at these times.

We set out to assess preferences for cotton nesting material and to determine if material composition influences retrieval. Because of this, the project centered on the mouse-tick tube interface and led to other key observations regarding mouse collection behavior regarding efficiency of collection and risk exposure of prolonged retrieval. Further investigation is needed to understand permethrin-treated material utilization in nests and the efficacy of mouse treatment with acaricides before we can effectively predict and refine control strategies. Moreover, additional experimentation under different conditions may reveal ways to improve deployment for more effective use.

## Acknowledgments

We would like to thank the staff at the staff at the UW Arboretum, especially ecologist Brad Herrick. Jordan Mandli was supported by the Parasitology and Vector Biology Training Program T32AI007414. Sydney Ring was an undergraduate student supported by the Welton Sophomore Apprenticeship. This work was supported by research funding provided by Thermacell Repellents, Inc.

## VIDEOS

Videos are available as supplemental files submitted with manuscript or can be viewed online. Online database:

https://uwmadison.box.com/s/h4f35on5fsndy5mcut5gfkgurip3xryf

Clip 1: Two mice captured at same time

https://uwmadison.box.com/s/mur93bj9e1bi5na4kxlm9q9oayzrypmb

Clip 2: Rapid cotton ball removal

https://uwmadison.box.com/s/l1a0c8dukh57vdq418v9g2lrveiwgqcz

Clip 3: Removal of entire tube contents

https://uwmadison.box.com/s/dcry93gsmsnkya7t47b404fsi9515lgk

Clip 4: Cotton batting removal

https://uwmadison.box.com/s/liemap3x0e8v2guwzr8yu4vvqbyh5324

Clip 5: Cotton ball removal

https://uwmadison.box.com/s/z459ccb4gyy7k401k0n05ikxktcexa43

Clip 6: Double cotton ball removal

https://uwmadison.box.com/s/itstr44kri8d21kry8crpo0p1o2rvlag

Clip 7: Cotton ball and cotton batting removal

https://uwmadison.box.com/s/31ix5jixtpmb41corx3m52d3fckosqkj

